# Perinatal maternal chronic exposure to dibutyl phthalate promotes visceral obesity in adult female offspring

**DOI:** 10.1101/2021.07.27.454002

**Authors:** Kunyan Zhou, Ran Cheng, Meina Yang, Xiaoyang Shen, Xiaoyan Luo, Li Ma, Liangzhi Xu, Jing Zhang

**Affiliations:** Department of Obstetrics and Gynecology, West China Second University Hospital, Chengdu, Sichuan, People’s Republic of China; Reproductive Endocrinology and Regulation Laboratory, West China Second University Hospital, Sichuan, People’s Republic of China; Key Laboratory of Birth Defects and Related Diseases of Women and Children (Sichuan University), Ministry of Education, China; Surgery Department, West China Second University Hospital, Chengdu, Sichuan, People’s Republic of China

**Keywords:** dibutyl phthalate, DBP, obesity, visceral fat, glucolipid metabolism, PI3K, PTEN

## Abstract

**Introduction:** Maternal exposure to dibutyl phthalate (DBP) may result in glucolipid dysfunction in female offspring. However, the underlying mechanisms remain elusive. We hypothesized that chronic maternal DBP exposure could induce abnormal metabolism of glucolipid.

**Materials and methods:** Sprague-Dawley rats were intraperitoneally injected with different doses of DBP, estradiol, and corn oil from gestational day 7 until the end of lactation. The weights, visceral fat percentage, serum lipid, insulin and glucose, protein levels of PI3K signal pathway in muscle were detected in F1 female offspring.

**Results:** Although the birth weight of F1 female offspring was not different among groups, the weights were heavier in DBP groups from postnatal day 7 to adult (P<0.001). The visceral adipose percentage in adult female offspring was increased by perinatal exposure to DBP (P<0.001). Decreased serum levels of triglyceride (P<0.0001), fasting glucose (P=0.004), prolactin (P=0.006), HOMA-IR (P=0.014) were found in female offspring exposed to DBP, but no difference for fasting insulin, total cholesterol, adiponectin. Increased protein levels of p-AKT, but decreased PTEN and GPR30 were observed in muscle of female offspring in DBP group, but without significant difference. None difference was observed for the protein levels of PI3K, AKT, GLUT4, InsR and IRS-1.

**Conclusion:** Maternal perinatal exposure to DBP induced obesity and accumulation of visceral adipose tissue for the adult female offspring. Serum glucolipid and local signal transduction of PTEN/PI3K/AKT pathway in muscle were not adversely affected by perinatal exposure to DBP for adult female offspring.

## Introduction

Endocrine-disrupting chemicals (EDCs) are exogenous compounds which can alter hormone biosynthesis, causing adverse effects to human health and their offspring (Diamanti-Kandarakis et al., 2009). Phthalate esters is a kind of EDCs, commonly used in products such as plastics, medical equipment, personal care products, and in the coating of some oral medications. Phthalate metabolites have been detected in various body fluids including blood, urine, and follicular fluid (Du et al., 2016). Multigenerational and transgenerational effects on female health are induced by prenatal exposure to the phthalate mixture, such as metabolic syndrome (Manikkam et al., 2013). However, the long-term metabolic impacts of early-life phthalate and phthalate mixture exposures are controversial. Furthermore, the metabolic impacts of phthalate exposures have focused on diethylhexyl phthalate (DEHP) (Neier et al., 2019).

Dibutyl phthalate (DBP) is also a widely used plasticizer, is rapidly absorbed, distributed, and metabolized to mono-butyl-phthalate (Rodriguez-Sosa et al., 2014). DBP has been studied intensively on the reproductive toxicology in male development before. Interestingly, women of reproductive age have the highest exposure levels of mono-butyl-phthalate than any other age/sex group (Blount et al., 2000; Guo et al., 2011). The estimated daily intake in humans is 7-10μg/kg/day in the general population and 233μg/kg/day in patients taking DBP-coated medications (Hines et al., 2011). DBP metabolites can reach the ovary and have been measured in the follicular fluid of women (Du et al., 2016).

Obesity is associated with the development of insulin resistance, which in turn plays an important role in the obesity-associated cardiometabolic complications (Barazzoni et al., 2018). Recent systematic review suggested that insulin resistance was positively associated with DBP exposure, however, the association of obesity and phthalate exposure was unclear (Radke et al., 2019). The PTEN/PI3K/Akt signaling cascades have effect on glucose uptake via translocation of GLUT-4 (Li et al., 2017). This signal pathway can also play a role on the efficacy of EDCs on female (Hu et al., 2018).

Perinatal maternal exposure to DBP has been reported to influence the health of female offspring, but the results were controversial. Furthermore, the underlying molecular mechanisms of metabolic dysfunction induced by perinatal maternal exposure to DBP have not been clearly elucidated to date. In this study, we aimed to study the effects of perinatal DBP exposure on glucolipid metabolism in female offspring and examine the role of PTEN/PI3K/AKT signaling pathway.

## Material and methods

### Animals and tissue sample

All experiments were conducted according to the Guide for the Care and Use of Laboratory Animals of National Research Council and approved by the Ethical Scientific Committee for the Care of Animals at West China Second University Hospital, Sichuan University (No. 2018-007).

Adult female and male Sprague-Dawley rats were purchased and allowed to acclimate to the facility for two weeks before use. The rats were maintained in polysulfone cages at Animal Facility of West China Second University Hospital, Sichuan University, under controlled conditions (22 ± 1°C, 12h light/dark cycle). Food and water were provided for ad libitum consumption. Groups of two females were mated with one male overnight and the day of the vaginal plug was considered day 0 of gestation. Pregnant females were housed individually with hard wood shavings as bedding and randomly allocated into five groups using a random number table.

Pregnant rats were treated intraperitoneally (i.p.) with DBP (99.5% pure, solid, Sigma Chemical Co., St. Louis, MO, USA) in corn oil (Sigma, 8001-30-7) at the doses of 33 mg/kg/day, 66 mg/kg/day, 132 mg/kg/day from gestational day 7 (GD7) throughout post-natal day 21 (PND 21); with estradiol (E_2_, 20 μg/kg/day) or corn oil (negative group, 0.3 ml/day) as controls. The administration dosage and route of DBP was chosen based on the previous study describing transgenerational inheritance of reproductive disease with perinatal exposure to DBP (Manikkam et al., 2013). Reproductive disorder has also been proved by our unpublished preliminary study with the same administration method of DBP. The weights of gestational F0 rats were weighted every week during gestation and lactation (PND 21).

At weaning, only female F1 offspring were selected and housed in five groups with three individuals per cage, including around 10 cages in each group. The weights of F1 offspring were weighted regularly up to adult age (PND 90). Then F1 females were anesthetized with isoflurane and killed by carbon dioxide overdose. The muscles or white visceral fat tissue were collected, weighted and stored at −80°C until use. The percentage of visceral fat tissue was calculated by dividing fat weight by body weight. Fasting blood samples were collected from the heart and separated by centrifugation at 3000 rpm for 15 min at 4 °C. Serum samples were stored at ≤ −80°C. Total cholesterol (Tch), triglyceride (TG), fasting glucose were measured using biochemical detection kits (KHB, Shanghai, China) in automatic biochemical analyser (Hitachi 7600).

### Enzyme-linked immunosorbent assay

Levels of prolactin (PRL), fasting insulin (FINS), adiponectin (ADP) in the serum were measured in duplicate using enzyme-linked immunosorbent assay (ELISA) kits (Jiancheng Nanjing, China), according to the manufacturer’s instructions. The detection limits were 0.5-200 ng/ml, 1-300mIU/L, 0.1-30mg/L for prolactin, insulin and adiponectin, respectively. The ELISA kit intra-assay coefficient of variation and inter-assay coefficient of variation were < 10% and < 12%, respectively. Homeostatic Model Assessment of estimated Insulin Resistance (HOMA-IR) was calculated based on fasting glucose and FINS (So et al., 2020).

### Western blot analysis

Antibody against β-actin, insulin receptor (InsR), insulin receptor substrate-1 (IRS-1) and HRP-conjugated secondary antibodies were obtained from Zen-Bioscience Company (Chengdu, Sichuan, China); antibody against phosphatidylinositol 3-hydroxy kinase (PI3K), Phosphorylated protein kinase B (p-AKT), and glucose transporter 4 (GLUT4) from Proteintech Group Inc. (Wuhan, China); antibody against AKT, phosphatase and tensin homology deleted on chromosome 10 (PTEN), G protein coupled estrogen receptor 30 (GPR30) were from abclonal Technology.

Muscle was lysed using RIPA Lysis Buffer (Beyotime Inst Biotech, Shanghai, China). The bicinchoninic acid (BCA) protein assay kit (Pierce, IL, USA) was used to determine the protein concentrations. 20-40 μg of lysate protein was resolved by 10% SDS-PAGE. Proteins were transferred to 0.45 μM polyvinylidene fluoride (PVDF) membranes after electrophoresis. The membranes were incubated with 5% nonfat milk for 90 min at 37°C, then washed with TBST and incubated at 37°C with primary antibodies for 2 hours. After a thorough wash with TBST and incubation with respective HRP-conjugated secondary antibodies at 37°C for 1 hour, the membranes were incubated for 2-5min in enhanced chemiluminescence reagent (Bio-Rad) and were then exposed to film for signal detection. The optical density of target protein was corrected using β-actin and analyzed with Image J.

### Statistical analysis

All data were presented as the mean and standard deviation (SD). Analysis of variance (One-way) was used to conduct multiple comparisons between groups and was then followed by Bonferroni post hoc comparisons if equal variances were assumed or Dunnett’s T3 post hoc comparisons if equal variances were not assumed. Statistical analyses were performed using SPSS v.20 software (IBM, Inc.). A P-value < 0.05 was statistically significant.

## Results

### The weights change of F0 and F1 rats

The basal weights of F0 pregnant rats were not different among groups at the stage of GD0 and GD7. However, after the DBP was injected, the gain weight of F0 rats was fewer in DBP group than that of control group from GD14 to PND21 (P<0.001), especially in the medium and high DBP groups (Table 1).

**Table 1.** The weight of F0 rats among groups during pregnant and lactation period.

| Days | Negative control<br>(n=8) | Low DBP<br>(n=9) | Medium DBP<br>(n=9) | High DPB<br>(n=7) | *P* |
| --- | --- | --- | --- | --- | --- |
| GD0 | 244.3±4.80 | 242.1±1.54 | 243.8±1.72 | 243.1±3.72 | 0.53 |
| GD7 | 265.9±8.86 | 260.2±2.77 | 263.7±1.58 | 261.6±5.88 | 0.17 |
| GD14 | 290.13±9.65 | 279.0±3.81 | 284.8±2.90 | 278.4±5.88 | < 0.001 |
| GD21 | 317.1±12.56 | 298.3±7.23 | 299.0±5.43 | 293.6±6.08 | < 0.001 |
| PND0 | 289.6±5.60 | 284.3±3.20 | 282.2±3.15 | 281.4±3.73 | < 0.001 |
| PND7 | 293.1±4.87 | 292.6±6.06 | 285.1±3.14 | 280.8±3.13 | < 0.001 |
| PND14 | 291.1±4.12 | 287.1±4.86 | 280.0±3.81 | 274.4±3.21 | < 0.001 |
| PND21 | 283.4±7.27 | 278.3±4.36 | 274.1±4.19 | 271.3±1.98 | < 0.001 |

DBP, dibutyl phthalate; GD, gestational day; PND, post-natal day; SD, standard deviation

The birth wights of F1 female rats in DBP groups were not different from that of control (P>0.05). However, the weights began growing heavier in DBP groups than that of control from PND 7 until adult (PND90), especially heavier in the high dosage group of DBP (Table 2).

**Table 2.**
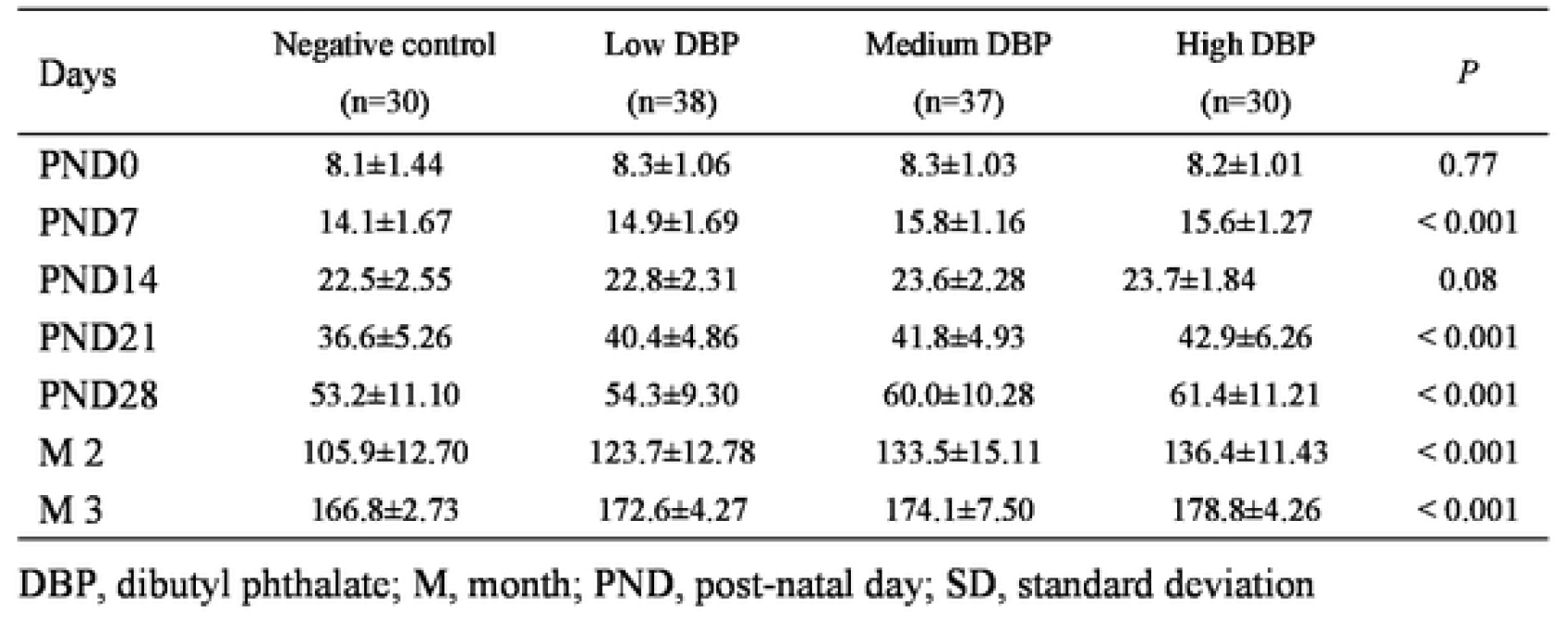
The weights of F1 rats among groups.

| Days | Negative control<br>(n=30) | Low DBP<br>(n=38) | Medium DBP<br>(n=37) | High DBP<br>(n=30) | *P* |
| --- | --- | --- | --- | --- | --- |
| PND0 | 8.1±1.44 | 8.3±1.06 | 8.3±1.03 | 8.2±1.01 | 0.77 |
| PND7 | 14.1±1.67 | 14.9±1.69 | 15.8±1.16 | 15.6±1.27 | < 0.001 |
| PND14 | 22.5±2.55 | 22.8±2.31 | 23.6±2.28 | 23.7±1.84 | 0.08 |
| PND21 | 36.6±5.26 | 40.4±4.86 | 41.8±4.93 | 42.9±6.26 | < 0.001 |
| PND28 | 53.2±11.10 | 54.3±9.30 | 60.0±10.28 | 61.4±11.21 | < 0.001 |
| M 2 | 105.9±12.70 | 123.7±12.78 | 133.5±15.11 | 136.4±11.43 | < 0.001 |
| M 3 | 166.8±2.73 | 172.6±4.27 | 174.1±7.50 | 178.8±4.26 | < 0.001 |

DBP, dibutyl phthalate; M, month; PND, post-natal day; SD, standard deviation

### The glycolipid metabolism of F1 female offspring

The visceral adipose tissue was much heavier in DBP group than that of control with the growing up of offspring. The percentage of visceral fat was much higher in DBP group for adult female offspring, especially in the medium and high DBP groups (Table 3). Being consistent with the effects of E_2_, the serum levels of TG (P<0.0001), fasting glucose(P=0.004), PRL (P=0.006), HOMA-IR (P=0.014) in F1 female offspring were decreased after perinatal exposure to DBP (Fig 1). However, the levels of FINS, Tch, ADP were not different among groups.

**Table 3.**
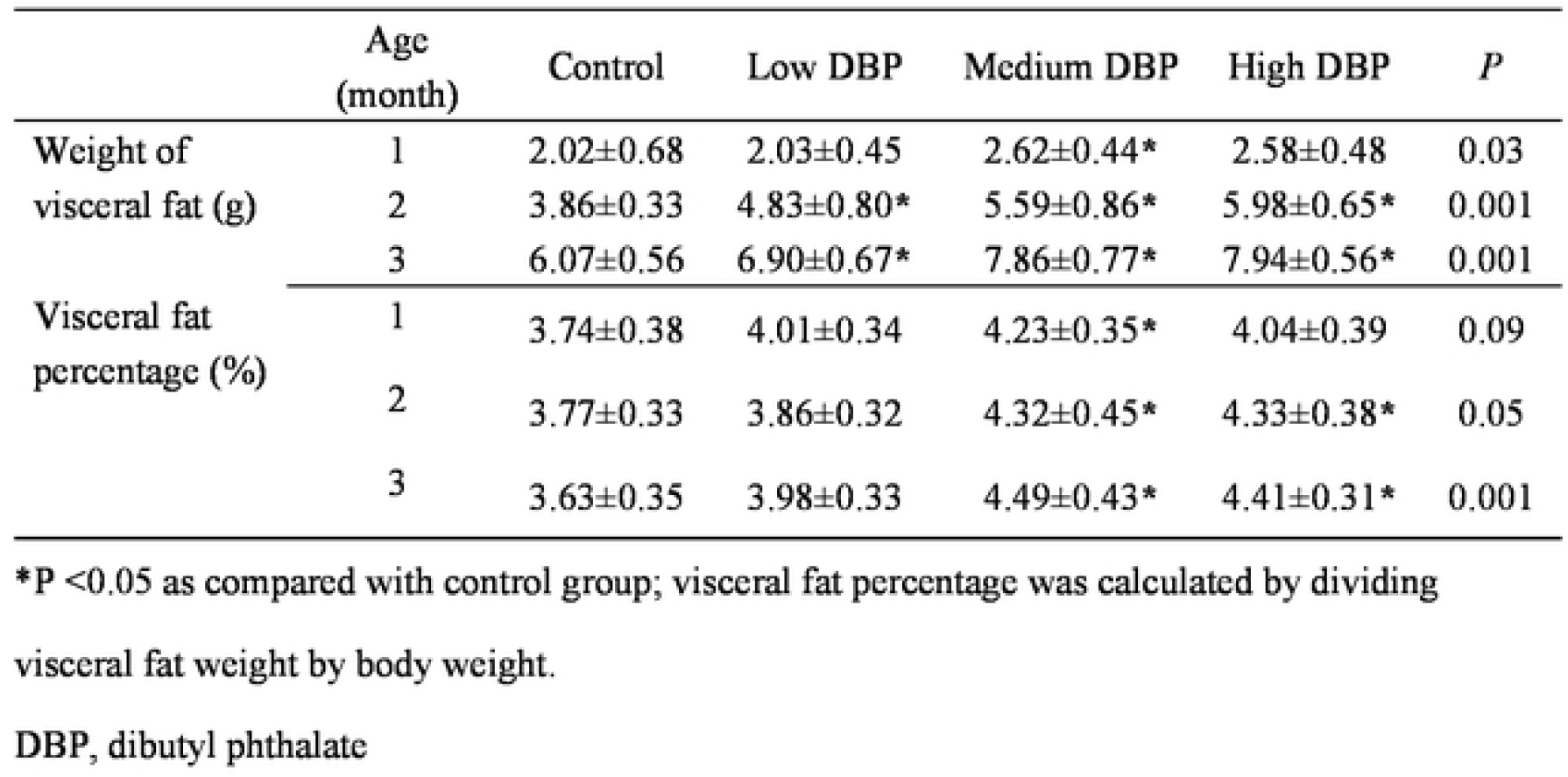
The visceral fat of F1 female rats in different age among groups.

|  | Age<br>(month) | Control | Low DBP | Medium DBP | High DBP | *P* |
| --- | --- | --- | --- | --- | --- | --- |
| Weight of<br>visceral fat (g) | 1 | 2.02±0.68 | 2.03±0.45 | 2.62±0.44* | 2.58±0.48 | 0.03 |
|  | 2 | 3.86±0.33 | 4.83±0.80* | 5.59±0.86* | 5.98±0.65* | 0.001 |
|  | 3 | 6.07±0.56 | 6.90±0.67* | 7.86±0.77* | 7.94±0.56* | 0.001 |
| Visceral fat<br>percentage (%) | 1 | 3.74±0.38 | 4.01±0.34 | 4.23±0.35* | 4.04±0.39 | 0.09 |
|  | 2 | 3.77±0.33 | 3.86±0.32 | 4.32±0.45* | 4.33±0.38* | 0.05 |
|  | 3 | 3.63±0.35 | 3.98±0.33 | 4.49±0.43* | 4.41±0.31* | 0.001 |

\*P <0.05 as compared with control group; visceral fat percentage was calculated by dividing visceral fat weight by body weight.
DBP, dibutyl phthalate

**Fig 1.**
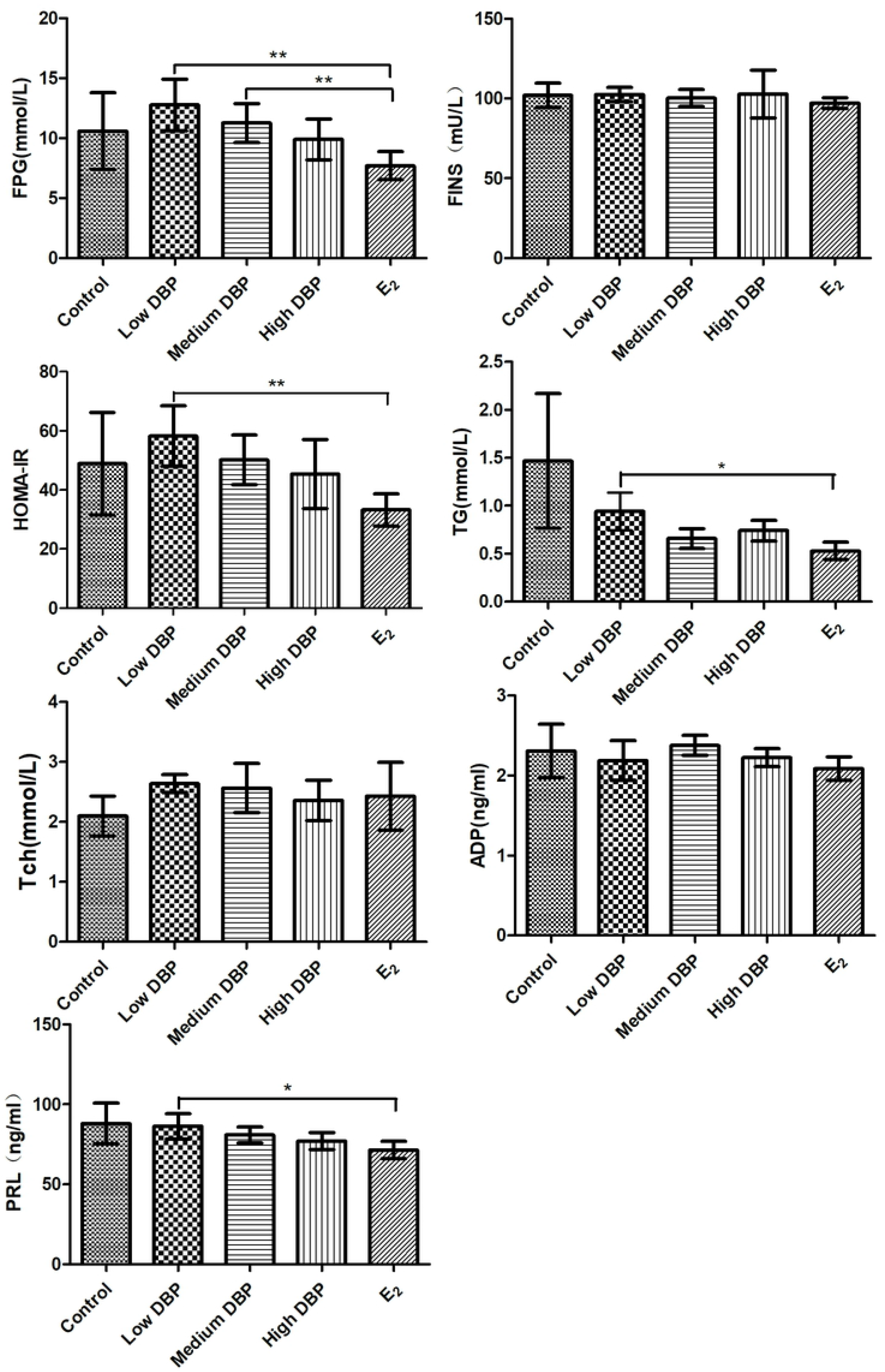
The serum levels of glucolipid in female offspring after the perinatal exposure to dibutyl phthalate (*P<0.05 and **P<0.01 in post hoc comparisons)

### The levels of proteins in glucolipid metabolism expressed in muscle

Increased p-AKT, but decreased PTEN and GPR30 were observed in muscle tissue of female offspring exposed to DBP, but without significant difference (Fig 2, Fig 3). However, the protein levels of PI3K, AKT, GLUT4, InsR, IRS-1 were not different among control and DBP groups. The protein levels of GLUT4, PTEN (p=0.029) and GPR30 (P=0.006) were significantly increased by E2.

**Fig 2.**
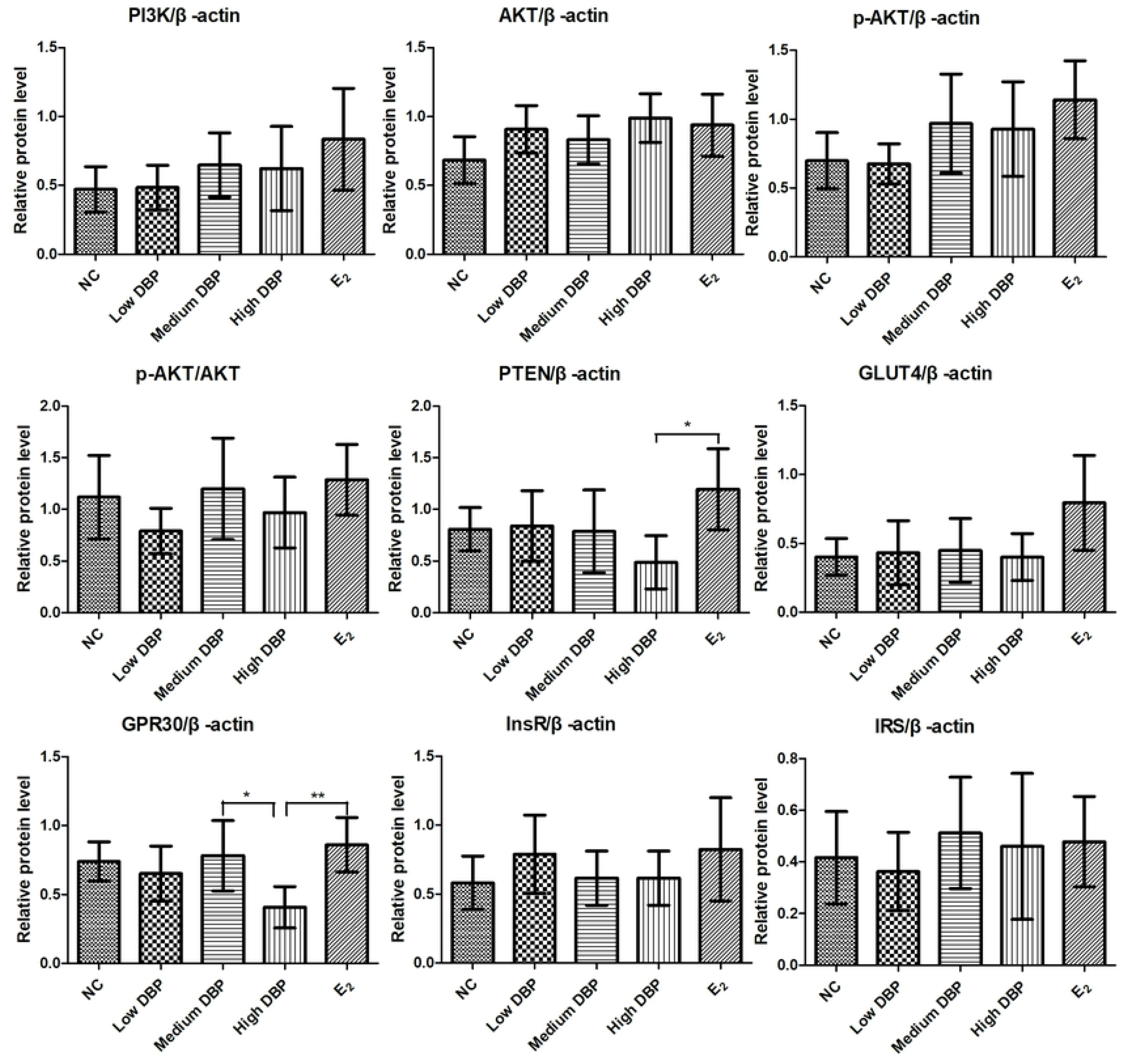
The levels of proteins in glucolipid metabolism expressed in muscle of female offspring after maternal perinatal exposure. Columns show the result of densitometric analysis, which is corrected and normalized by β-actin.

**Fig 3.**
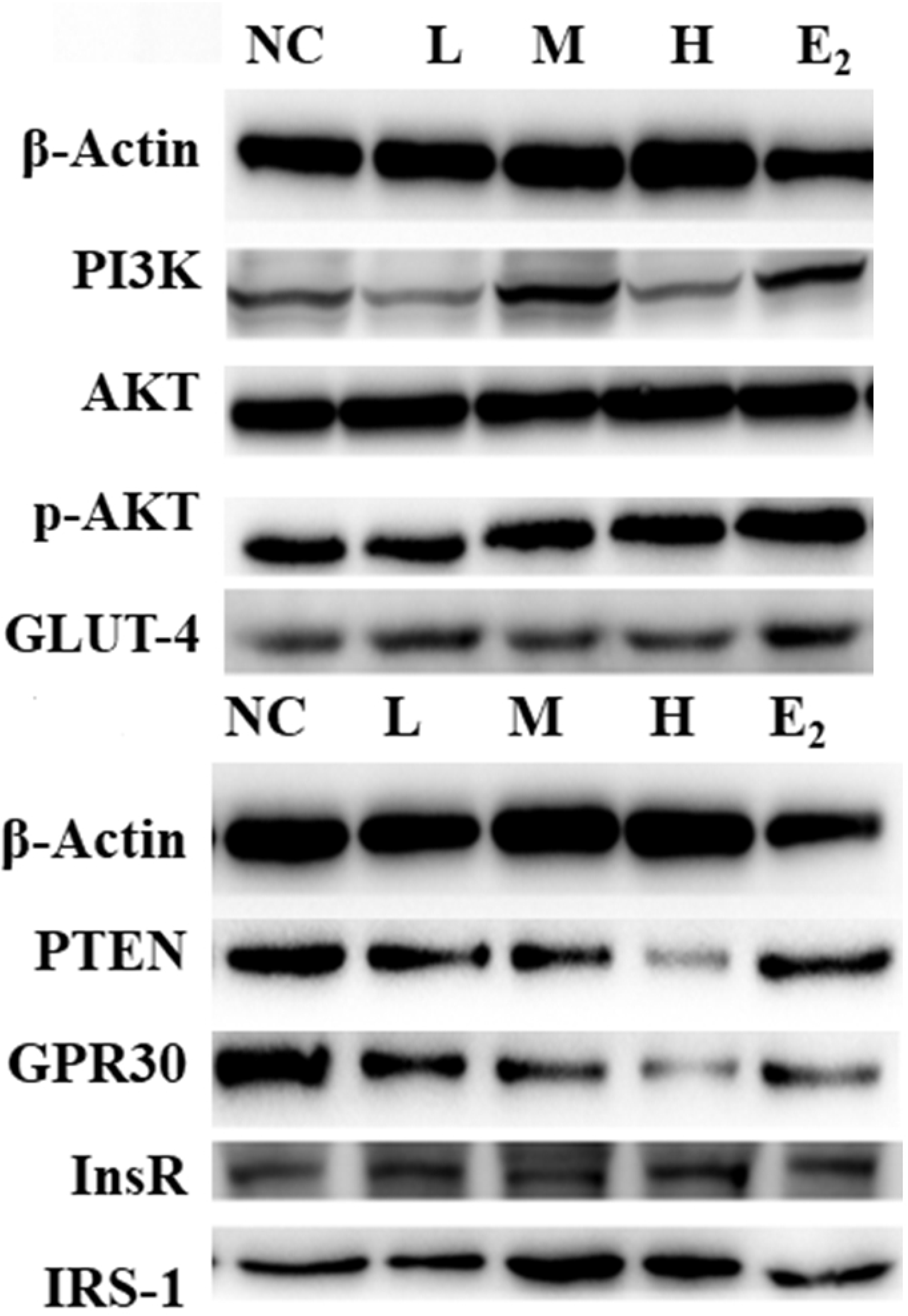
The levels of proteins in glucolipid metabolism expressed in muscle of female offspring evaluated by western blot.

## Discussion

In present study, the glucolipid metabolism and the underlying molecular mechanisms induced by the perinatal maternal exposure to DBP were explored. Decreased weight gain of F0 rats, but increased weight and visceral adipose accumulation of adult female offspring were found. Serum glucolipid metabolism and local signal transduction of PTEN/PI3K/AKT pathway in muscle tissue were not adversely affected by perinatal exposure to DBP for adult female offspring.

The mechanism and various phthalate effects on human glucose metabolism remain largely unknown. The metabolic impacts of developmental phthalate exposures have been focused on DEHP and found that females perinatally exposed to DEHP only had increased body fat percentage and decreased lean mass percentage, whereas females perinatally exposed to diisononyl phthalate (DINP) only had impaired glucose tolerance (Neier et al., 2019). The metabolic impact of phthalate is discrepant among different components; the impact of phthalate mixtures is also different from that of single phthalate. The efficacy of DBP might be different from those of DEHP or DINP.

In present study, heavier adult weight, and higher percentage of visceral fat tissue were found in adult female offspring perinatally exposed to DBP. Decreased serum levels of triglyceride, fasting glucose, prolactin, HOMA-IR were also found in female offspring, but no difference for fasting insulin, total cholesterol, and adiponectin.

Being consistent with the results of present study, intrauterine exposure to low-dose DBP (5 mg/kg/day) from GD 12 until PND 7 also promoted obesity in adult female and male offspring, but with evidence of glucose and lipid metabolic disorders and a decreased metabolic rate (Li et al., 2020). Xiong et al. (2020) also found the body weight of mice was increased after exposure to both low and high doses of DBP (0.1 and 1 mg/kg) by oral gavage. However, the serum levels of hepatic triglyceride and total cholesterol were increased in this study. Moreover, disturbed homeostasis of gut microbiota, hepatic lipid metabolism disorder, liver inflammation were also found in mice exposure to DBP. Consistently, sub-chronic exposure to low concentration of DBP increased body weight gain, feed efficiency, abdominal to thoracic circumference ratio, and body mass index in rats. Meanwhile, serum cholesterol decreased, glucose increased with DBP treatments (Majeed et al., 2017). However, exposure to 50mg/kg/day DBP alone induced a marked decrease in insulin secretion and glucose intolerance, but had no influence on insulin resistance (Deng et al., 2018).

From the results of above studies, the weight was identified to be increased when exposed to DBP. However, the effects of DBP exposure on lipid and glucose metabolism was controversial. The result discrepancy among the studies might be induced by the different administration route, duration, period, and dosage of DBP. The transgenerational effects of DBP on F1 offspring might be different from that of F0 generation. The function of liver and gut might be severely disturbed by the oral route, which could furtherly adversely affect the glucolipid metabolism.

The effective mechanism of DBP on glucolipid metabolism remain largely unknown. Fundamental research suggests that DBP contamination accelerate glucose consumption and upregulate the expression of porins and periplasmic monosaccharide ATP-binding cassette transporter solute-binding proteins for the metabolism of sugars in microbes (Chen et al., 2020). DBP-containing food or feeding adults DBP food affects the expression of homologous genes involved in xenobiotic and lipid metabolism (Williams et al., 2016). In present animal study in vivo, increased protein of p-AKT, decreased PTEN and GPR30 were observed in muscle of female offspring in DBP group, but without significant difference. None difference was observed for the protein levels of PI3K, AKT, GLUT4, InsR and IRS-1 by us.

Cell experiments in vitro suggest the combined effect of DEHP and DBP promotes a ROS-mediated PI3K/Akt/Bcl-2 pathway-induced pancreatic β cell apoptosis that is significantly higher than the effects of each PAE (Li et al., 2021). Wang et al. found that the expression of PTEN protein was higher, while the expression of p-PI3K1, p-AKT, p70S6K and 4E-BP1 protein in the PI3K/AKT/mTOR signal pathway were significantly decreased in DBP-induced apoptosis of testicular Sertoli cells in rats (Wang et al., 2017).

Risk assessment case study indicate that DBP-induced downregulation of genes in the lipid/sterol/cholesterol transport pathway as well as effects on immediate early gene/growth/differentiation, transcription, peroxisome proliferator-activated receptor (PPAR) signaling and apoptosis pathways in the testis (Euling et al., 2013). Perinatal phthalate exposures are associated with short- and long-term activation of PPAR target genes in liver tissue, which manifested as increased fatty acid production in early postnatal life and increased fatty acid oxidation in adulthood (Neier et al., 2020). Intrauterine exposure of mice to low-dose DBP (5 mg/kg/day) appears to promote obesity in offspring by inhibiting UCP1 via endoplasmic reticulum stress (higher expression of Bip and Chop), a process that is largely reversed by treatment with TUDCA (Li et al., 2020). DBP aggravate type 2 diabetes by disrupting the insulin signaling pathway and impairing insulin secretion. DBP exposure could disrupt the PI3K expression and AKT phosphorylation, and decrease the level of GLUT-2 in the pancreas tissue (Deng et al., 2018).

Mechanistic studies have characterized the mode of action for DBP in the glucolipid metabolism using muscle, liver, pancreas, testis and adipose tissue. The discrepancy among the above results may be attributed to the type of experiment (in vivo vs. in vitro), route of DBP administration (orally, intraperitoneally, vs. addition to the medium), dosage of DBP (low or high), duration of exposure (acute or chronic; single or complex), different tissues (liver, muscle, pancreas, testis or adipose tissue).

The strength of present study is to explore the long-term metabolic consequence of perinatal chronic maternal exposure to DBP for the adult female offspring, to explore the environmental deleterious effects on developmental origins of adult metabolic dysfunction. The limitation of present study is that the administration route (i.p.) of DBP is not the common environmental exposure mode (oral or skin exposure). Furthermore, given the widespread exposure of humans to numerous contaminants, the combinatorial effects of multiple chemicals also merit evaluation.

## Conclusion

Maternal perinatal exposure to DBP could induce visceral obesity for the adult female offspring. Serum glucolipid and local signal transduction of PTEN/PI3K/AKT pathway in muscle are not adversely affected by perinatal exposure to DBP for adult female offspring.

## Declarations

### Consent for publication

Not applicable

### Availability of data and materials

The data and materials could be available by contacting the corresponding author upon reasonable request.

### Authors’ contributions

ZJ: Conceptualization, writing original draft, investigation, data curation and analysis, project administration, funding acquisition. CR: Investigation, Data curation and analysis. YM, SX, LX, ML: Investigation, validation. XL: Resources, supervision. ZK: Conceptualization, methodology, validation, funding acquisition. All authors read and approved the final manuscript

## Acknowledgments

The authors thank Professor Bin Zhou for sample preservation; Siyu Zhou and Sicong Li for sample collection.

